# Human Genome Assembly in 100 Minutes

**DOI:** 10.1101/705616

**Authors:** Chen-Shan Chin, Asif Khalak

**Affiliations:** Foundation for Biological Data Science

## Abstract

De novo genome assembly provides comprehensive, unbiased genomic information and makes it possible to gain insight into new DNA sequences not present in reference genomes. Many de novo human genomes have been published in the last few years, leveraging a combination of inexpensive short-read and single-molecule long-read technologies. As long-read DNA sequencers become more prevalent, the computational burden of generating assemblies persists as a critical factor. The most common approach to long-read assembly, using an overlap-layout-consensus (OLC) paradigm, requires all-to-all read comparisons, which quadratically scales in computational complexity with the number of reads. We assert that recently achievements in sequencing technology (i.e. with accuracy ~99% and read length ~10-15k) enables a fundamentally better strategy for OLC that is effectively linear rather than quadratic. Our genome assembly implementation, Peregrine uses <u>**s**</u>parse <u>**hi**</u>erarchical <u>**m**</u>ini<u>**m**</u>iz<u>**er**</u>s (SHIMMER) to index reads thereby avoiding the need for an all-to-all read comparison step. Peregrine can assemble 30x human PacBio CCS read datasets in less than 30 CPU hours and around 100 wall-clock minutes to a high contiguity assembly (N50 > 20Mb). The continued advance of sequencing technologies coupled with the Peregrine assembler enables routine generation of human de novo assemblies. This will allow for population scale measurements of more comprehensive genomic variations -- beyond SNPs and small indels -- as well as novel applications requiring rapid access to de novo assemblies.

## Introduction

The initial human genome project and the development of technologies of cheap DNA sequencing technologies have advanced both academic research and industrialization of using genomic information to improve human health^1,2^. The fast decreasing cost of second generation sequencing technologies makes population scale study of certain type of variations, e.g., SNPs and small indel variants, possible. It leads to valuable information for genotype and phenotype association and many important and clinical relevant applications^3–5^. Meanwhile, the recent development long read technologies, e.g. sequencers from Oxford Nanopore Technology (ONT) and Pacific Biosciences (PacBio), can read DNA sequences that are orders of magnitude longer than those of second generation sequencing. With longer read lengths, it makes de novo assembly relatively easier and we can generate more contiguous assemblies^6–13^. The de novo reconstruction of a genome reduces the dependence on using a reference as prior information. A re-sequencing approach depending on a reference may not be effective to explore those genomic structures that are deviated from the references significantly. Recent studies with directly human genome assemblies have discovered new sequences that are not in the current reference genome^12,14–16^. Systematic approach for discovering structural variations identifies many new structural variations and projects that we still needs more samples to generate a more comprehensive catalogs of larger variants in human population other than SNPs and small indels.

The current barrier for big scale utilization of long reads is the relative cost and throughput compared to the second generation sequencing technologies. We expect the long read sequencing cost will follow the trajectory of the second generation sequencing which will continue to decrease. On reducing the computing cost of genome assembly, there are also many recent progresses that significantly reduce the total amount computational resources needed^17,18^. In a recent report, researchers were able to assemble 30x human genome in a couple hundreds to thousands of CPU hours with consensus reads that were longer than 10kb with average accuracy better than 99%^8,19^.

For long-read genome assembly, overlap-layout-consensus (OLC) paradigm^20,21^ is used in most of the current long-read assemblers for production. The quadratic comparison between reads remains the main bottleneck for further improving computation efficiency. For example, the hierarchical genome assembly process, HGAP^10^, initially designed for assembly noisy PacBio reads takes two overlapping steps, one for error correction and one for assembly graph generation needs 20,000 to 30,000 cpu hours to assemble a human genome from noisy sequences. Most high performance assemblers developed recently, e.g. Flye^17^, wtdbg2^18^ and Shasta^22^, adapt new strategies to avoid such expensive explicit overlapping steps between two full reads. Such optimization is likely necessary for efficiently assembling noisy long reads. Meanwhile, we can start with consensus reads with better accuracy to improve computation efficiency of overlapping reads for genome assembly. We find it is possible to reduce the computation complexity of the overlap detection by exploiting the better read accuracy. We have developed a new genome assembler Peregrine (https://github.com/cschin/peregrine). It utilizes an efficient indexing scheme to reduce computation time as low as 20 cpu hours for assembling a human size genome starting with consensus reads.

The genome assembly approach presented here can effectively make genome assembly become routine work without special setup for cluster computation. Simplifying such fundamental process for a genome project is key to routinely generating de novo genome assembly that avoids missing information from re-sequencing approach. The cost of de novo approach is currently more expensive than re-sequencing. Nevertheless, with the fast pace in advance of computational methods and sequencing technologies, the cost to generate de novo human genome may drop to a price point that it has become affordable for personalized medicine soon. Our method will also help to build pan-human-genomics references which allows us to capture novel human genome sequences that are not available in the current human reference GRCh38^12,15^. Given the potential to provide more comprehensive view for human genomes, we are hoping, the whole genome assembly approach will provide important information for genetic diseases that can not be revealed easily with re-sequencing approach.

## Results

We developed a new method for indexing sequence reads (See Figure 1 and Methods) to identify overlaps between two reads and implement a new genome assembler “Peregrine” with it. We test the Peregrine assembler on a number of public human genome datasets with different parameter sets. The full summary of the results in terms of computational resource usage are in Table 1. We utilized large memory compute nodes, e.g. Amazon Web Services *m5d.metal* (384 Gb RAM) or *r5d. 12xlarge* (384 Gb RAM) instance types, so we can run 24 indexing and overlapping processes concurrently. Peregrine uses 9 to 25 cpu core hours depending the sequencing coverage and parameters. The wall clock time of a typical assembly run ranges from one to two hours from initial fasta or fastq files to final assembly. The overlap-to-assembly graph module in Peregrine currently runs on a single core in about 30 to 40 minutes of wall clock time, depending on sequencing coverage and read length. This step takes a significant fraction of the overall wall clock time. It may be possible to further optimize this step with a parallel computing approach in the future. The single overlapping process may use memory up to 120Gb RAM depending on the sequencing coverage and read lengths. When running the overlapping processes concurrently the sequence data used for alignment confirmation are shared among concurrent processes through a memory mapped file approach^23^. When we split the full index into 24 partitions, each overlap process requires about ~ 8 to 10Gb memory for ~30x human genome sequencing to store the index. The design for index partitioning is simple and flexible in Peregrine. In order to run Peregrine in a computer with smaller memory, we can just increase the number of partitions and run the overlapping process in serial.

**Figure 1.**
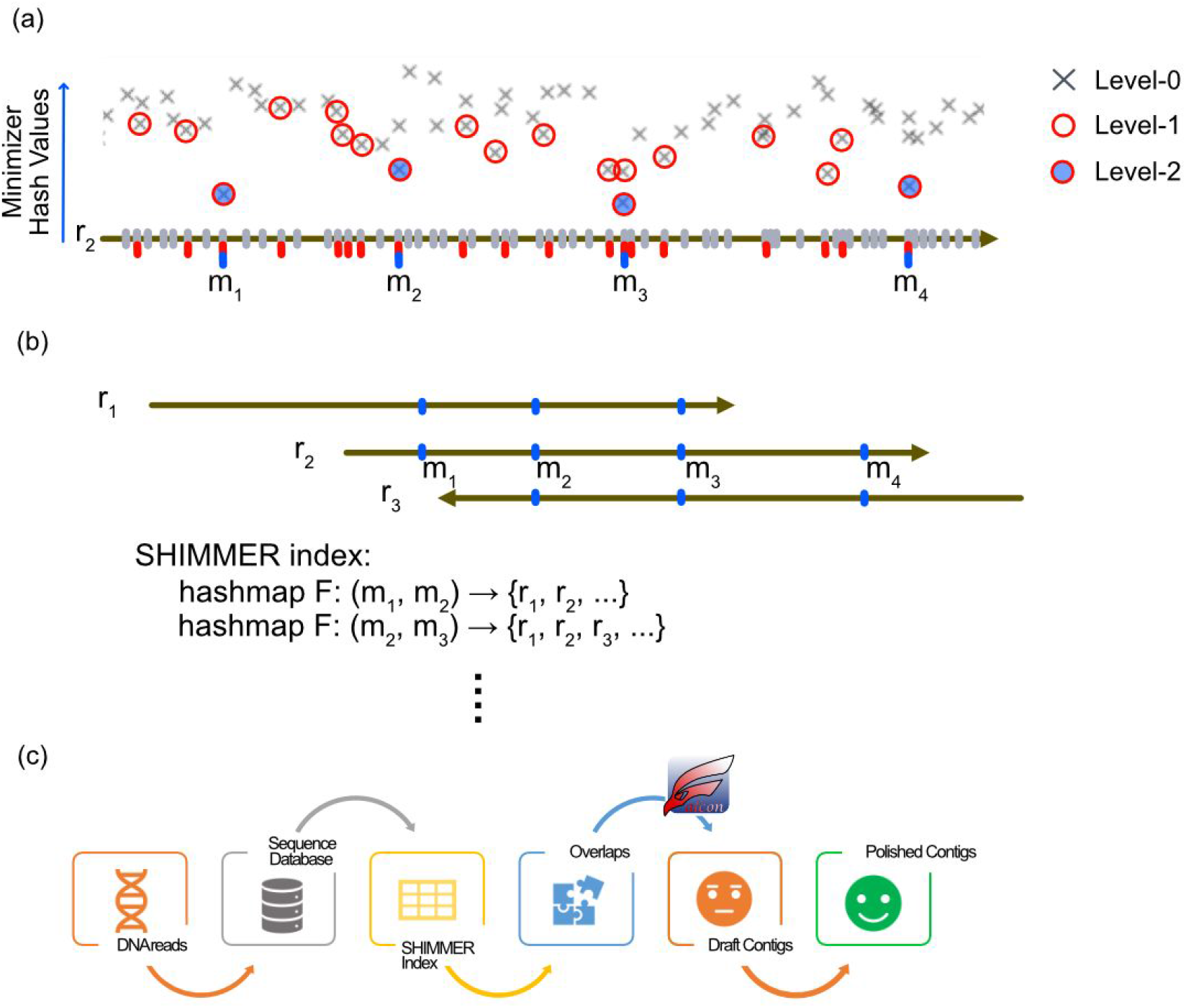
SHIMMER index generation and the Peregrine assembler workflow. (a) The gray tick-marks represent the locations of the level-0 minimizer along a read. The crosses represents the hash value of the minimizers. The level-1 minimizers (red tick-marks and circles) are the local minima of the windows through the neighboring minimizers. Similarly, the level-2 minimizers (blue tick-marks m_1_ to m_4_, and filled circles) are local minima of the level-1 minimizers over moving windows. (b) For each read, we scan the level-2 minimizers and generate a hash map that maps neighboring minimizer pair to a set of reads to speed up overlap finding. (c) The Peregrine assembler workflow. The overlapping module of Peregrine generates file that is compatible to FALCON assembler’s overlap-to-contig modules. After we get the draft contigs from FALCON assembler, we apply the FALCON-sense algorithm to polish the draft contig to increase the contig accuracy.

**Table 1(a).**

| Disk Usage |  |  |  |  |  |  |  |
| --- | --- | --- | --- | --- | --- | --- | --- |
| Genome | parameters | read length | # reads | seqdb | L0 index | L2 or L1 index | Peregrine version |
| HG002 | r=3, l=2 | 13478.3 | 6610240 | 89.1 Gb | 44.7 Gb | 15.2 Gb | 0.1.5.3 |
| HG002 | r=4, l=2 | 13478.3 | 6610240 | 89.1 Gb | 44.7 Gb | 9.51 Gb | 0.1.4.rc8 |
| HG002 | r=6, l=2 | 13478.3 | 6610240 | 89.1 Gb | 44.7 Gb | 4.46 Gb | 0.1.4.rc4 |
| HG002 | r=18, l=1 | 13478.3 | 6610240 | 89.1 Gb | 44.7Gb | 4.84 Gb | 0.1.4.rc6 |
| HG002 | r=36, l=1 | 13478.3 | 6610240 | 89.1 Gb | 44.7Gb | 2.38 Gb | 0.1.4.rc6 |
| HG002 (Sequel II) | r=6, l=2 | 11301.3 | 3895357 | 44.2 Gb | 24.2 Gb | 5.13 Gb | 0.1.4.rc4 |
| chm13 | r=4, l=2 | 10960.1 | 6902752 | 75.7 Gb | 38.6 Gb | 8.20 Gb | 0.1.4.rc4 |
| chm13 | r=3, l=2 | 10960.1 | 6902752 | 75.7 Gb | 38.6 Gb | 13.08 Gb | 0.1.4.rc4 |
| NA12878 | r=3, l=2 | 9963.2 | 9082038 | 90.5 Gb | 45.3 Gb | 15.34 Gb | 0.1.5.0 |

**Table 1(b).**
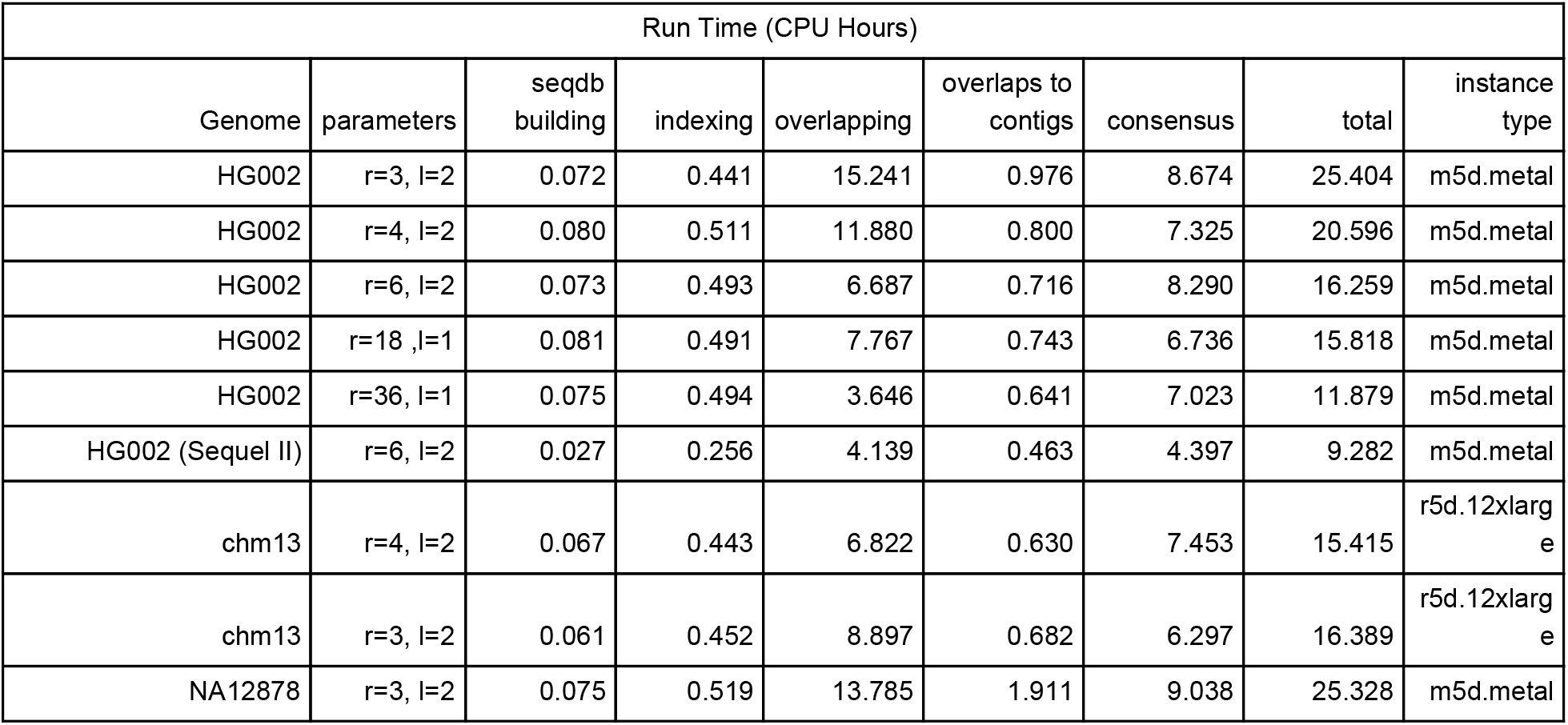

| Run Time (CPU Hours) |  |  |  |  |  |  |  |  |
| --- | --- | --- | --- | --- | --- | --- | --- | --- |
| Genome | parameters | seqdb building | indexing | overlapping | overlaps to contigs | consensus | total | instance type |
| HG002 | r=3, l=2 | 0.072 | 0.441 | 15.241 | 0.976 | 8.674 | 25.404 | m5d.metal |
| HG002 | r=4, l=2 | 0.080 | 0.511 | 11.880 | 0.800 | 7.325 | 20.596 | m5d.metal |
| HG002 | r=6, l=2 | 0.073 | 0.493 | 6.687 | 0.716 | 8.290 | 16.259 | m5d.metal |
| HG002 | r=18, l=1 | 0.081 | 0.491 | 7.767 | 0.743 | 6.736 | 15.818 | m5d.metal |
| HG002 | r=36, l=1 | 0.075 | 0.494 | 3.646 | 0.641 | 7.023 | 11.879 | m5d.metal |
| HG002 (Sequel II) | r=6, l=2 | 0.027 | 0.256 | 4.139 | 0.463 | 4.397 | 9.282 | m5d.metal |
| chm13 | r=4, l=2 | 0.067 | 0.443 | 6.822 | 0.630 | 7.453 | 15.415 | r5d.12xlarge |
| chm13 | r=3, l=2 | 0.061 | 0.452 | 8.897 | 0.682 | 6.297 | 16.389 | r5d.12xlarge |
| NA12878 | r=3, l=2 | 0.075 | 0.519 | 13.785 | 1.911 | 9.038 | 25.328 | m5d.metal |

**Table 1(c).**

| Assembly Statistics Summary |  |  |  |  |  |
| --- | --- | --- | --- | --- | --- |
| Genome | parameters | Total | N50 | max | # seq |
| HG002 | r=3, l=2 | 2,915,831,858 | 35,323,946 | 108,396,683 | 2,571 |
| HG002 | r=4, l=2 | 2,906,581,718 | 33,364,927 | 108,396,082 | 2,438 |
| HG002 | r=6, l=2 | 2,896,918,528 | 27,848,727 | 108,387,980 | 2,398 |
| HG002 | r=18, l=1 | 2,897,055,690 | 26,459,768 | 102,052,462 | 2,384 |
| HG002 | r=36, l=1 | 2,894,701,861 | 6,746,698 | 32,267,778 | 3,194 |
| HG002 (Sequel II) | r=6, l=2 | 2,872,604,008 | 21,936,975 | 109,829,231 | 2,022 |
| chm13 | r=4, l=2 | 2,839,105,926 | 29,260,433 | 102,007,116 | 978 |
| chm13 | r=3, l=2 | 2,838,663,153 | 33,307,555 | 95,435,791 | 844 |
| NA12878 | r=3, l=2 | 2,880,983,308 | 25,483,577 | 109,846,404 | 3,061 |

The overlapping module generated a file that is compatible with the one used in the FALCON assembler^24^. After the overlapping step, Peregrine uses the overlap-to-contig layout modules from the FALCON assembler to generate contigs. The accuracy of overlap detection can affect the contig lengths. Given our contig N50 lengths are on par with those generated by FALCON with daligner, we expect the performance of detecting the correct overalps is close to the daligner for these accurate reads. However, the Peregrine overlapper is designed specifically to find overlaps that are longer than a couple hundred to thousand bases. Small overlaps induced by repeats that is not significant for constructing contigs properly will be ignored by the design of the overlapping searching algorithm.

To evaluate Peregrine’s overlapping module, we simulate reads using E. coli genome and test the overlapping performance at different level of error rates and length distributions. For average 15kb reads and 1% error, within 55,935 overlapping read pairs detected, 99.94% are true positive and there 32 false positives which are caused by repeats in the genome. We are able to detect 53,858 (99.77%) of 53,982 true overlap pairs with overlap length > 10,000 bp. When the error rates become higher, the false negative rates can become higher. Using more dense or multiple indexes may help to detect overlaps with higher error rates. For larger and more complicated genomes, false positive rates can increase because more repetitive sequences in the genome. Even with the false positives and false negatives, the Peregrine assembler generate one contig with 100% identity to the E. coli reference sequence from the simulated read set.

The major improvement of Peregrine from FALCON for long low-noisy sequences is (1) speeding up for read-to-read overlap detection and (2) polishing the draft-contig through consensus to increase the contig base accuracy. The draft contig generated by Peregrine will have the same error rate of the input sequences. Peregrine maps the reads back to the draft contig and apply an updated FALCONsense algorithm ^25^ to polish the draft contig.

We perform the Benchmarking Universal Single-Copy Orthologs (BUSCO) evaluation with the vertebrata lineage profile^26^ on the selected assemblies of four different human genomes. The BUSCO completeness ranges from 93.8% to 95.2% (Table 2). For this BUSCO evaluation, the Peregrine’s results are on-par or higher than most recently reported de novo human genome assemblies from similar data^19^. Our CPU core hour usage is significantly lower than other assemblers previously applied to the same HG002 dataset and achieve the same or slightly better BUSCO performance. For assembly contiguity, our results are also on-par or better than those reported previously.

**Table 2.**

|  | parameters | Complete (%) | Complete Single Copy (%) | Complete and duplicated (%) | Fragmented (%) | Missing (%) | Total BUSCO groups |
| --- | --- | --- | --- | --- | --- | --- | --- |
| HG002 | r=3, l=2 | 95.0 | 93.7 | 1.3 | 2.3 | 2.7 | 2586 |
| CHM13 | r=3, l=2 | 94.0 | 92.8 | 1.2 | 2.7 | 3.3 | 2586 |
| NA12878 | r=3, l=2 | 95.2 | 93.9 | 1.3 | 2.3 | 2.5 | 2586 |

For accuracy assessment for the Peregrine’s consensus polishing module, we utilize the orthogonal sequenced VMRC59 BAC sequences collected for the hydatidiform mole human genome CHM13^8^. Voller and colleagues identified 31 BACs that are not intersected with segmental duplication (SD) regions for assessing the assembly accuracy (Figure 2(a)). For the 31 non-SD BACs, the estimated error rates in Phred QV scale range from 25 to 52 with a mean at 42.2. Out of 4.64Mb from the 31 BACs that are fully aligned to the assembly contigs, there are only 1 mismatch and the rest of the errors are from 337 insertions and 604 deletions. For other 310 VMRC59 BACs which may come from segmental duplicated regions, only 78 are fully aligned and the estimated error rates are significant highly than the non-SD regions (Figure 2(b)). The long repeats from those SD regions are definitely challenges for the assembler’s overlapping and contig layout models.

**Figure 2.**
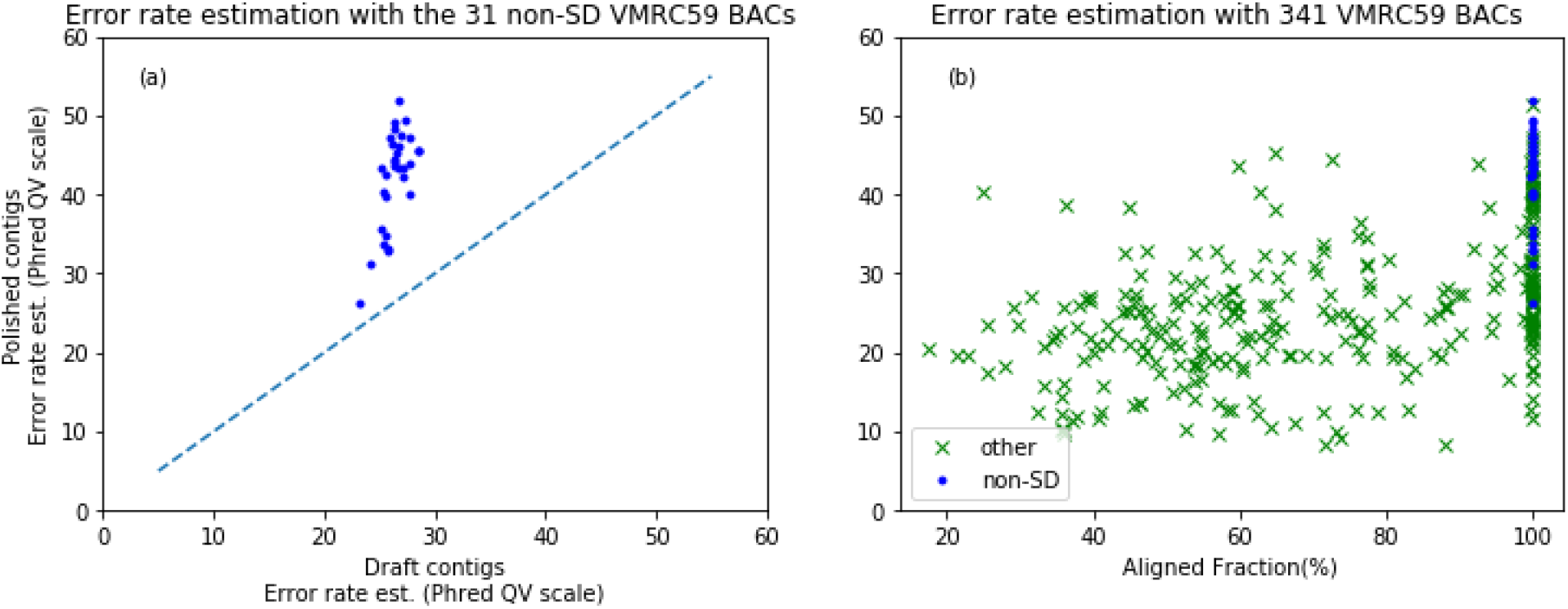
Error rate estimation using VMRC59 BACs and the CHM13 assembly (a) Error estimation comparison for draft and polished contigs using the 31 non-SD BACs. All data point is above the y=x dashed line showing the consensus module significantly reduces the error rates. (b) The estimated errors vs. the aligned fraction for all 341 VMCRC59 BACs.

## Methods

### Indexing Reads With Sparse Hierarchical Minimizer

A minimizer (Yorke 2004) is a k-mer that is one of a curated list of *k*-mers such that any significant overlapping exact match between reads is composed of *w* consecutive *k*-mers contained in the list. Such lists, which characterize significant matches will generically be *far* smaller in size than the reads themselves. The process of generating minimizers is akin to database indexing. We extend this concept further from a list of minimizers to a hierarchy of lists of minimizers. Given a list of lower level minimizers, we can identify a hierarchical subset of minimizers to further reducing the index size.

In detail, the level-0 minimizers are the k-mers which have lowest hash values over the moving windows along the read sequence. The hash function is usually chosen to avoid picking minimizers from simple context, e.g. homopolymer stretches. The details of the hash function are typically not important, other than avoiding collisions over the set of k-mers. After generating the level-0 minimizers, we scan through the list of level-0 minimizers and identify the subset of minimizers which themselves have the lowest hash values in the level-0 list over moving windows of a given size. We call this new subset of minimizer level-1 minimizers and the size of the window as reduction factor *r*. Similarly, we can repeat the same process over the level-1 minimizers to create a hierarchical structure of minimizers. In our implementation, we generate one or two extra levels of minimizers from the level-0 ones for indexing the reads.

An example of the process generating different levels of minimizers over a simulated read with 1% error from E. coli is shown in Figure 3. We retain the original hash values and the positions of the minimizers in the reads in the different level of indices. In the simulated E. coli dataset, the index size of level-2 minimizers is just 10% (4.98 Mbyte) of the level-0 minimizer index (49.9 M byte) for window size *w*=80, kmer size *k*=16 and reduction factor *r*=6 between the level.

**Figure 3.**
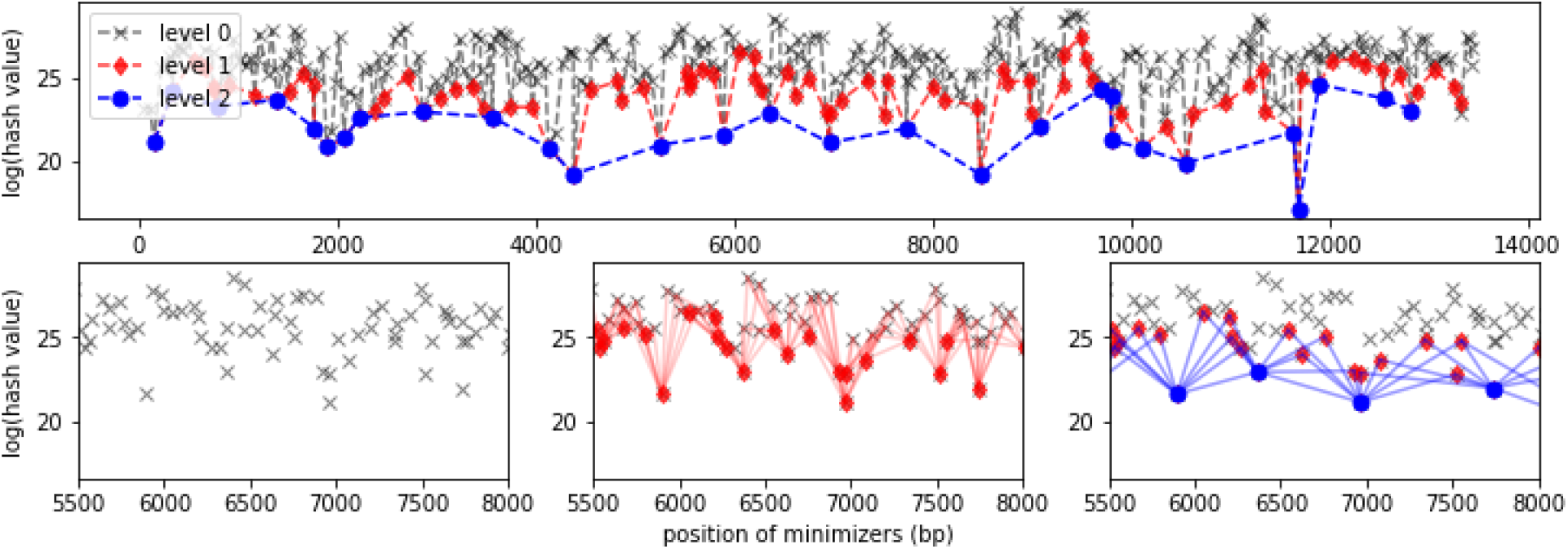
Example of two level of minimizer reductions for a single read. Upper Panel: The locations and hash values of multiple level minimizers across a read. The black crosses are the level 0 minimizers compute with k=16 and w=80. The red diamonds and the blue dots are the level-1 and level-2 minimizer respectively. Lower Panel: The steps for building higher level SHIMMERs. Left: the level-0 minimizer in the given window. Middle: The level 1 minimizers (red diamonds) are the minimizers from those level 0 minimizers. The red lines connect the level 1 minimizers to those level 0 ones that the level 1 minimizers are derived from. Right: the level 2 minimizers (blue dots) are generated by finding the minimizers of the level 1 minimizers.

### Aggregating Reads by Minimizer Pairs and Confirming The Overlaps

We build a hashmap using neighboring minimizer pairs (NMP) as the key from the last level of minimizer list of each read. The value of the hashmap of a NMP is the list of read identifier where NMP can be identified in those reads. Each NMP can be considered as a digest that represents the sequence context across their span. For example, the distance between two neighboring level-0 minimizers is 47.3 +/− 1.8 bp with *w*=80 and *k*=16 in the E. coli dataset. In such case, the two sequences of 47 bps with the same NMP will has likelihood that they are identical to each other. The NMP from higher level minimizers spans larger greater range across the reads. The distance between two neighboring level-2 minimizers is 527.9 +/ 74.6 bp after two levels of reduction of the minimizer with a reduction factor r=6 (Figure 4). It is about 7.1 times longer than the neighboring distance of the level-0 minimizers. As sparse hierarchical minimizers are used for such indexing schema, we call this “SHIMMER index”.

**Figure 4.**
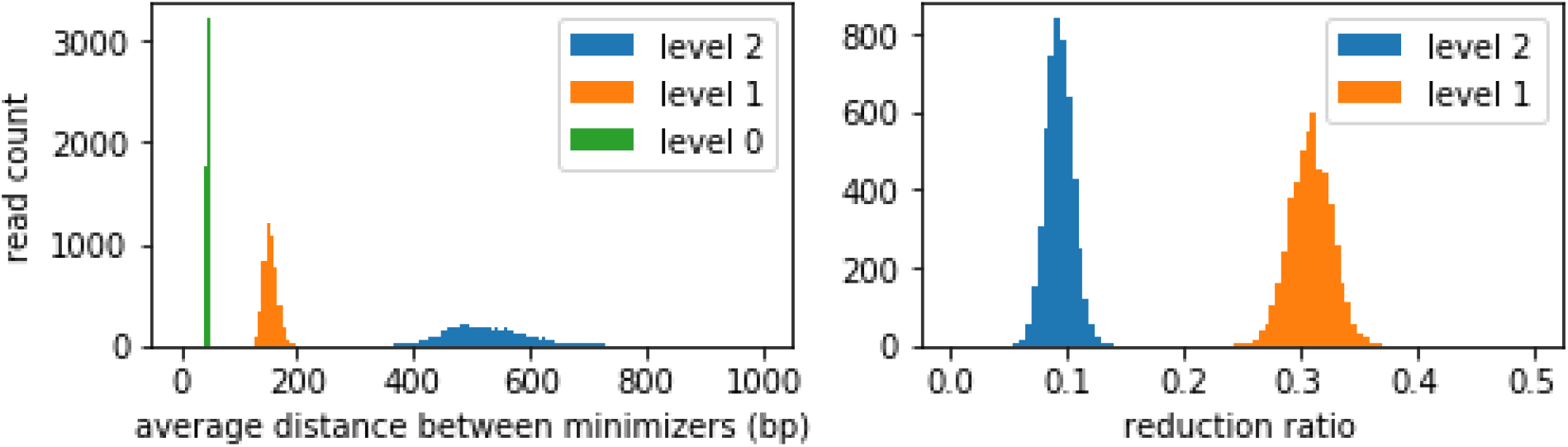
Left: The distribution of the average distance between minimizers of different level. Right: the distribution of the reduction ratio (= (number of level 1 or 2 minimizers) / (number of level 0 minimizers) per read).

Figure 5 shows the number of unique NMP keys and the number of reads that are gathered by each NMP. To avoid indexing NMP with two minimizers that are too close to each other, we impose a minimum distance requirement. Two minimizers that are within 100bp are not used for indexing. As the most level-1 or level-2 minimizers are spread out, this filter will have little effect on overall overlapping performance but it helps to avoid high density NMPs from a repetitive region getting indexed.

**Figure 5(a).**
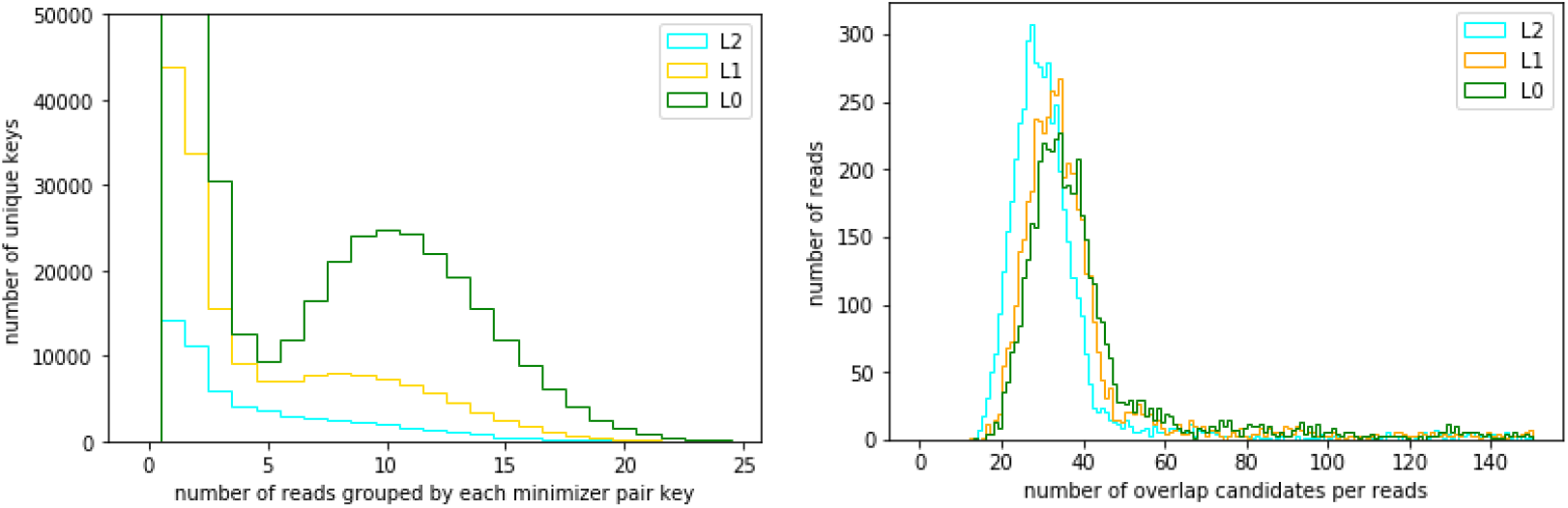
Without filtering the NMPs that the minimizers are close to each other: Left: the distribution of number of hits per unique NMP. Right: the distribution of the number reads that can be identified as overlapped reads using all NMPs in a read. (Average number of hits per NMP key: level-0: 483869 unique NMPs, 6.15 hits / per key, level-1: 174744 unique NMPs, 5.19 hits / per NMP, level-2: 57842 unique NMPs, 4.68 hits / per NMP. Average Number of read overlaps from each read: level-0: 51.3, level-1: 44.8, level-2: 32.5.)

**Figure 5(b).**
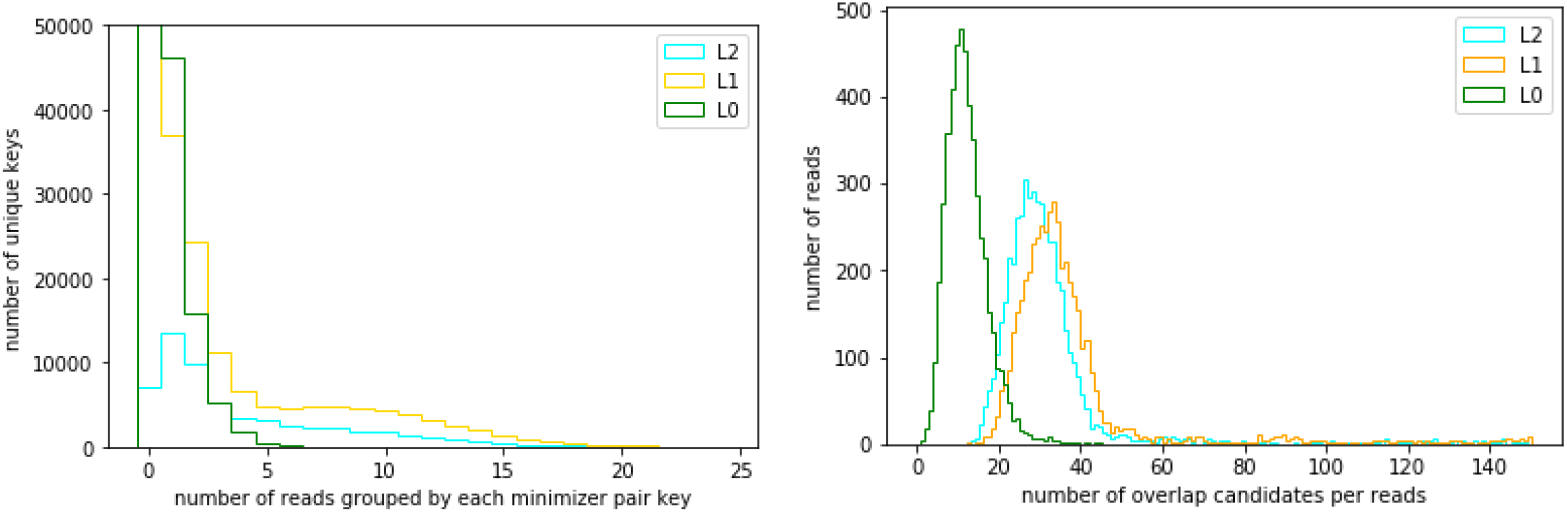
With a filter removing NMPs that the minimizers are less than 100bp apart: Left: the distribution of number of hits per unique NMP. Right: the distribution of the number reads that can be identified as overlapped reads using all NMPs in a read. (Average number of hits per NMP key: level-0: 483869 unique NMPs, 0.218 hits / per key, level-1: 174744 unique NMPs, 3.24 hits / per NMP, level-2: 57842 unique NMPs, 3.95 hits / per NMP. Average Number of read overlaps from each read: level-0: 11.67, level-1: 42.8, level-2: 31.9.)

The reads that are grouped by each NMP has high probability coming from the same region of the genome and we perform a detailed alignment^27^ between the reads of each group to confirm the overlaps. As we only need to compare the reads with each group, we avoid global read-to-read comparison. Multiple NMPs may identify the same read pairs for overlapping, we record reads that has been tested for overlapping to avoid duplicated computation.

Long repetitive sequences in genome usually pose computational challenges for the overlapping step. For example, if a sequence are repeated *M* times and the sequencing coverage is *C*, there are potentially *M*^2^*C*^2^ reads are mostly identical to each other. The extra *M*^2^ factor can significantly increase the computation time and storage requirements for the read overlapping step. Various heuristics are used to reduce this extra computation burden due to repeats^18,25,28^. In our indexing schema, if an NMP is mapped to too many reads, we know the sequence that the NMP represents are from repetitive sequences in the genome, and we choose to ignore them for the initial assembly and process them with extra steps (Supplementary Figure S1).

### Generating Consensus

The SHIMMER indexing schema are also useful for fast read to contig or contig to reference mapping. For the consensus step, we map the reads back to the contig using the SHIMMER indices from the contigs and the reads. Once the reads are mapped to each contig is then used to generate the final consensus to improve the contig accuracy.

The final consensus base accuracy is affected by multiple factors: (1) input quality in terms of read length, accuracy and overall sequencing coverage, (2) repeat content and (3) heterozygosity. For simulated *E. coli* reads, we are able to achieve 100% accuracy after consensus as the errors in the simulated reads are mathematically “random” and lend themselves to straightforward denoising approaches. Real world sequencing data (especially for eukaryotes) are rarely sufficiently random for straightforward denoising, both in read errors and in sequence content. On the heterozygous SNP sites, the algorithm is designed to pick an allele with majority read support and may fail to pick the correct one if there are complicated multiple local variants between the two haplotypes. Our current assembly process after the SHIMMER indexing and overlapping steps does consider diploid or polyploid genome. The techniques developed by earlier work. e.g. FALCON-Unzip^25^ or Canu Trio binning^29^, can be adapted to the downstream or the upstream processing to obtain pseudo-haplotigs or haplotigs.

## Discussion

The overlap-layout-consensus (OLC) approach is advantageous for de novo sequencing because the raw read information is largely preserved in the final assembly, making it possible to trace back to original reads that provide direct evidence of the content of contigs in the assembly. In previous work, this has only been possible by performing an all read-to-read comparison in the form of a string graph^20^ or an overlap graph^21^. This implies that the number of read comparisons scales quadratically with the number of input reads. It becomes the main computational bottleneck for the overlap-based assembly approach. Our proposal for hyper-rapid assembly (i.e. in 100 minutes) overcomes quadratic scaling with a linear pre-processing step.

Our method scans all the reads to construct a hashmap that records the read locations of neighboring minimizer pairs (NMPs), which act as indices in the hashmap. Let N be the total number of reads, C be the sequencing coverage and L be the average read length, the number of operations to build the hashmap is proportional to the total number bases ~ G*C* = NL, where *G* is the genome size. The number of hashmap indices is therefore proportional to the genome size *G*. For each unique hashmap index across the entire genome, the number of reads associated with this index is proportional to the sequencing coverage *C*. Thus, the algorithmic runtime complexity to construct the SHIMMER index is O(*GC*) or O(*NL*).

Assuming roughly uniform distribution across indices, applicable to non-repetitive sequence regions, the computational cost to determine overlaps scales quadratically with coverage, *C*. This is due to an all to all comparison within each subset of reads that shares an index - that is, for the same neighboring minimizer pair. The number of minimizer indexes in the index is *G/d*, where *d* is the average distance between the minimizers, about 500 (Figure 4). The overall runtime complexity for the comparison step is thus, O(*GC*^2^/*d*), which compares favorably with standard OLC approaches whose algorithmic runtime complexity is quadratic with the number of reads O(*N*^2^) or O(*G*^2^*C*^2^/*L*^2^), where each read must be compared against all others to checking the overlapping condition. The ratio of complexity estimates from standard OLC to SHIMMER, *Gd/L*^2^, is roughly 5000x for current reads on a human-size genome where *G* ~ 10^9^, *d* ~ 500, and *L*~ 10^4^. We anticipate that if *L* increases substantially, then adjustments to indexing parameters may be needed to increase *d* for optimal results. In practice, we similarly find dramatic benefits of SHIMMER indexing from a runtime perspective.

For any eukaryotic genome, one must identify repeats in the early stages of assembly so as to avoid unnecessary computation. In highly repetitive genome regions, the *C*^2^ scaling breaks down because a single NMP may correspond to many loci in the genome, each over which has a certain coverage. The NMPs that are likely to correspond to long repetitive sequences are therefore identified in the indexing process since the number of corresponding reads is greater, in a statistically significant way, from that predicted by uniform coverage. These receive special treatment in the current implementation, and reads which have no unique part in the genome will be ignored. However, it would be straightforward to include these reads in the final assembled contigs for further processing without compromising the runtime complexity.

The current SHIMMER indexing method has been designed and tested on reads longer than 10kb and with an error rate smaller than 1%. When the error rate increases from this, the sensitivity of finding correct minimizer pairs decreases. For instance, we might not find all overlapping pairs necessary for constructing the assembly graph to lay out contigs successfully. There is a natural dependency of the final genome assembly quality on the input sequencing read quality in terms of length and accuracy. It might be possible to adapt the current SHIMMER index approach to accommodate lower accuracy reads by modifying the indexing parameters and incorporating additional denoising steps to the process. Meanwhile, if sequencing technology significantly improves read lengths beyond the current technology limit (~15kb), the use of sparse minimizers to index reads will become an even more effective approach for de novo assembly.

The performance characteristics of this method brings de novo human genome assembly towards being rapid, affordable, and universally accessible. We have demonstrated that a human genome assembly can be performed in about two hours with cloud-accessible hardware, on a single node. Similarly, a dedicated desktop computer with sufficient physical memory (e.g. 2019 Mac Pro) can also perform this task without the need for a cluster computer setup, which avoids both software infrastructure requirements and the need for specialized skills in grid computing.

This universal access, coupled with the speedy advance of DNA sequencing technologies, will enable routine generation of reference grade genome assemblies - as rapidly as one or two days from sample collection to result. Thus, de novo sequencing has been seen as solely investigational and essentially an end in itself - to document and discover the unknown sequences. We foresee these new capabilities as enabling the use of full de novo sequences as part of more complex efforts which integrate sample collection, sequencing, and other steps towards a larger goal in the areas of industrial, agricultural, and medical biotechnology. Both the completeness and timeliness of such outputs have potential for novel use-cases. The rapid turnaround will make this useful for diagnostic or confirmatory measurements - including clinical or therapeutic efforts one day.

## Acknowledgements

We like to thank Eichler Lab for sharing the CHM13 dataset for us to test out Peregrine. Special thanks to Mitchell R. Vollger and Glennis A. Logsdon for providing the information about VMRC59 BAC dataset for evaluating accuracy of the CHM13 assembly. We also like to thank many unsung scientists and engineers at PacBio who have figured out how to make longer and more accurate reads possible.

## Data Source

HG002:

ftp://ftp-trace.ncbi.nlm.nih.gov/giab/ftp/data/AshkenazimTrio/HG002_NA24385_son/PacBio_CCS_15kb/

HG002 (Sequel II):

https://downloads.pacbcloud.com/public/dataset/HG002_SV_and_SNV_CCS/

CHM13: https://www.ncbi.nlm.nih.gov/sra/SRX5633451

NA12878: https://www.ncbi.nlm.nih.gov/bioproject/PRJNA540705

## Supplementary Figures

**Figure S1.**
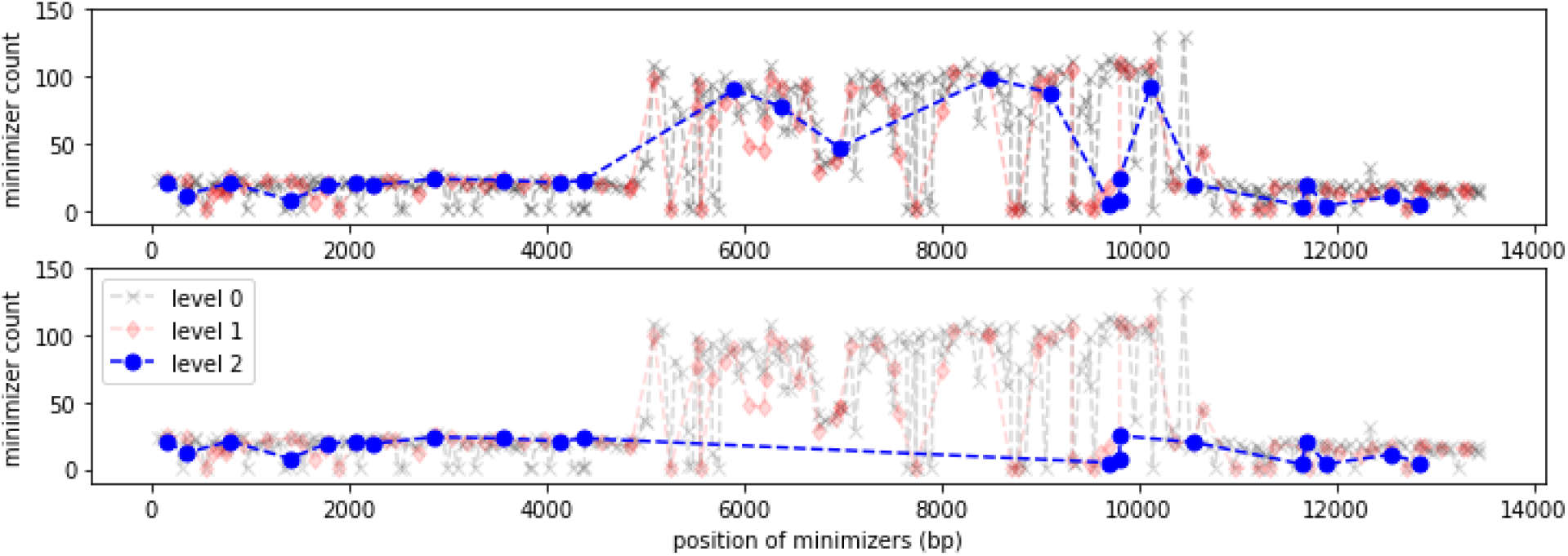
An example of filtering out high repeat count minimizers in a read. The minimizer count shows there is a repetitive segment from 5,000bp to 11,000bp in this read. If we filter out minimizers that has excessive count, e.g. greater than 50 in this example, we can reduce false hits caused by such repeats. We can still use the minimizers for the unique part to find proper overlaps. Upper: The blue dots show the level-2 minimizers before high repeat count filtering. Lower: The blue dots show the level-2 minimizers after filtering.

**Figure S2.**
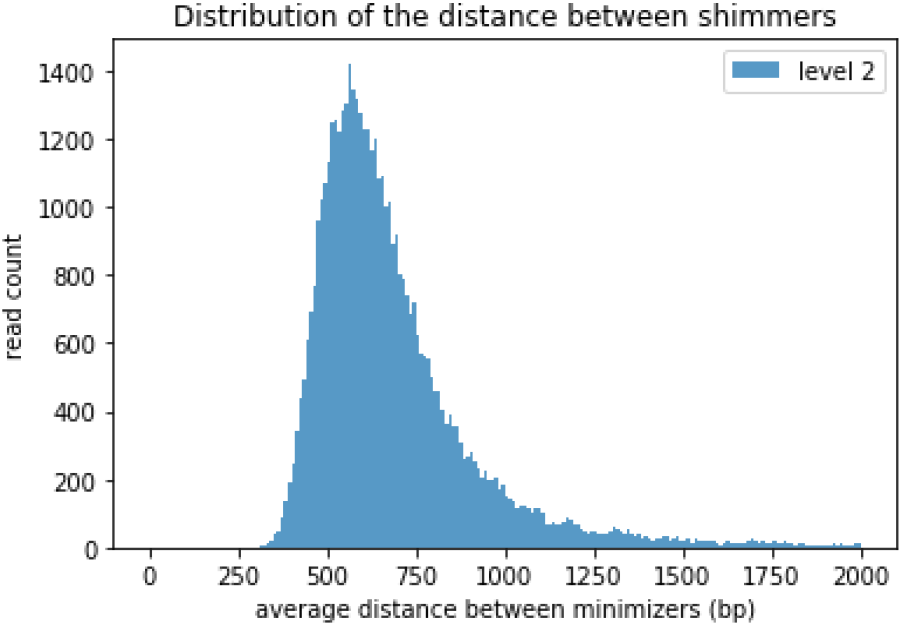
The distribution of the distances between level-2 minimizers (r=4) from a subset of reads in the 28x HG002 dataset. Right: The distribution of the number of overlap candidate identified per read from a subset of reads in the 28x HG002 dataset. It shows that we only need to check about 40 to 65 overlaps per read.

